# The BIR2/BIR3-interacting Phospholipase D gamma 1 negatively regulates immunity in Arabidopsis

**DOI:** 10.1101/815282

**Authors:** Maria A. Schlöffel, Andrea Salzer, Wei-Lin Wan, Ringo van Wijk, Maja Šemanjski, Efthymia Symeonidi, Peter Slaby, Joachim Kilian, Boris Maček, Teun Munnik, Andrea A. Gust

## Abstract

Plants have evolved effective strategies to defend themselves against pathogen invasion. Starting from the plasma membrane with the recognition of microbe-associated molecular patterns (MAMPs) via pattern recognition receptors, internal cellular signaling pathways are induced to ultimately fend off the attack. Phospholipase D (PLD) hydrolyzes membrane phospholipids to produce phosphatidic acid (PA), which has been proposed to play a second messenger role in immunity. The Arabidopsis PLD family consists of 12 members and for some a specific function in resistance towards a subset of pathogens has been shown. We demonstrate here that Arabidopsis PLDγ1, but not its close homologs PLDγ2 and PLDγ3, is specifically involved in plant immunity. Genetic inactivation of *PLDγ1* resulted in increased resistance towards the virulent bacterium *Pseudomonas syringae* pv. *tomato* DC3000 and the necrotrophic fungus *Botrytis cinerea*. As *pldγ1* mutant plants responded with elevated levels of reactive oxygen species to MAMP-treatment, a negative regulatory function for this PLD isoform is proposed. Importantly, PA levels in *pldγ1* mutants were not affected compared to stressed wild-type plants, suggesting that alterations in PA levels are unlikely the cause for the enhanced immunity in the *pldγ1* line. Instead, the plasma-membrane-attached PLDγ1 protein colocalized and associated with the receptor-like kinases BIR2 and BIR3, which are known negative regulators of pattern-triggered immunity. Moreover, complex formation of PLDγ1 and BIR2 was further promoted upon MAMP-treatment. Hence, we propose that PLDγ1 acts as a negative regulator of plant immune responses in complex with immunity-related proteins BIR2 and BIR3.

**One-sentence summary:** A phospholipase D is a novel negative regulator of plant immunity and forms complexes with regulatory receptor-like kinases.

## INTRODUCTION

Although plants are constantly in contact with potentially harmful microorganisms, disease development is rather an exception. This is due to an efficient plant surveillance system that detects highly conserved microbe-associated molecular patterns (MAMPs) via cell surface-located pattern recognition receptors to mount appropriate immune responses in a process called pattern-triggered immunity (PTI) (Couto and Zipfel, 2016; Saijo et al., 2018). One well studied example for such a MAMP is bacterial flagellin, of which a 22 amino acid long fragment is sufficient to trigger immune responses by binding to the extracellular leucine-rich repeat (LRR) domain of the receptor kinase FLAGELLIN SENSING 2 (FLS2) (Gómez-Gómez and Boller, 2000; Chinchilla et al., 2006). Upon ligand-receptor interaction, activated receptor complexes recruit co-receptors of the SOMATIC EMBRYOGENESIS RECEPTOR KINASE (SERK) family, most prominently BRASSINOSTEROID RECEPTOR1-ASSOCIATED KINASE1 (BAK1) (Chinchilla et al., 2007; Heese et al., 2007; Sun et al., 2013). In the absence of an immunogenic stimulus, however, the co-receptor kinase BAK1 is bound to the BAK1-INTERACTING RECEPTOR-LIKE KINASES BIR2 and BIR3 to avoid inappropriate receptor activation (Halter et al., 2014; Imkampe et al., 2017). Only upon ligand-binding to the corresponding immune receptor, BAK1 is released from BIR2/3 and can now join the receptor-ligand complex. Subsequently, cellular signaling events are triggered including the production of reactive oxygen species (ROS), the activation of mitogen-activated protein kinases (MAPKs) or the transcriptional reprogramming of the cell (Boller and Felix, 2009; Yu et al., 2017; Saijo et al., 2018) which ultimately leads to plant resistance against the microbial invader.

The activation of MAPKs and the oxidative burst can be modulated by lipid second messengers such as phosphatidic acid (PA) (Laxalt and Munnik, 2002; Testerink and Munnik, 2011), and PA levels have been shown to rapidly increase in various cell suspension cultures upon stimulation with different elicitors (van der Luit et al., 2000; Laxalt et al., 2001; den Hartog et al., 2003; de Jong et al., 2004; Bargmann et al., 2006). In plants, PA can be released from membrane phospholipids by two classes of enzymes, either directly via hydrolysis catalysed by phospholipase D (PLD) or via the combined action of phospholipase C (PLC) and diacylglycerol kinase (DGK) (Wang, 2004; Bargmann and Munnik, 2006; Arisz et al., 2009).

*Arabidopsis thaliana* (Arabidopsis) has 12 different genes for PLDs, and based on their gene architectures, sequence similarities, domain structures, and biochemical properties they have been grouped into the 6 subfamilies α, β, γ, δ, ε and ζ (Qin and Wang, 2002). PLDs play a role in lipid metabolism but are also involved in the regulation of cellular processes, such as hormone signaling, environmental stress responses and cellular and subcellular dynamics (Bargmann and Munnik, 2006; Wang et al., 2006). PLDα was one of the first PLD family members that have been characterised, and depletion of Arabidopsis *PLDα* resulted in decreased levels of PA and superoxide in leaf extracts (Sang et al., 2001). In guard cells, abscisic acid (ABA)-responses rely on the production of PA via PLDα1, and this PA is required for the accumulation of ROS by directly binding to and thereby stimulating NADPH oxidase activity (Zhang et al., 2009). Moreover, PLDα1 regulates the function of the heterotrimeric G protein GPA1, by interacting with the α-subunit and promoting the exchange of GTP to GDP, thereby mediating ABA-induced inhibition of stomatal opening (Mishra et al., 2006). A second member of this PLD subgroup, PLDα3, was also implicated in hyperosmotic stress and *pldα3* mutant plants were more sensitive to salinity and water deficiency (Hong et al., 2008). PLDs are also involved in plant growth as exemplified by PLDε of which mutant plants display decreased root growth and biomass accumulation (Hong et al., 2009).

In addition to the earlier observations in cell suspension cultures, PLDs have also been implicated in immunity of whole plants. Infection of Arabidopsis with the necrotrophic fungus *Botrytis cinerea* or the hemibiotrophic bacterial pathogen *Pseudomonas syringae* pv *tomato* (*Pto*) DC3000 resulted in increased PA levels, predominantly produced by PLDβ1 (Zhao et al., 2013). Genetic inactivation of *PLDβ1* lead to enhanced susceptibility to *Botrytis* infection, but increased resistance to *Pto*DC3000, accompanied by higher levels of ROS and salicylic acid (SA) (Zhao et al., 2013), indicating that PLDβ1 can have both positive and negative regulatory functions in plant defense, depending on the lifestyle of the pathogen. Specific PLD isoforms have also been shown to be important for penetration resistance, e.g. Arabidopsis *pldδ1* mutants fail to restrict penetration of the two powdery mildew fungi *Blumeria graminis* f. sp. *hordei* and *Erysiphe pisi* (Pinosa et al., 2013). In addition, the accumulation of chitin-induced defense genes appears to be delayed in the *pldδ1* mutant (Pinosa et al., 2013), suggesting a positive role for PLDδ1 in plant immunity.

We show here that Arabidopsis PLDγ1, a so far largely uncharacterized PLD isoform, plays a major role in immunity. Genetic inactivation of *PLDγ1* leads to increased resistance towards both bacterial and fungal infection, accompanied by elevated MAMP-induced ROS levels. However, alterations in PA levels are unlikely to be the cause for this increased resistance as they seem to be unaffected in *pldγ1* mutant plants. Rather, PLDγ1 can be found in complex with the BAK1-interacting LRR-RLKs BIR2 and BIR3, revealing a novel function for phospholipases in plant immunity.

## RESULTS

### PLDγ1 is the only PLDγ isoform involved in bacterial resistance

Whereas several Arabidopsis phospholipase D family members have been shown to be involved in plant immunity (Pinosa et al., 2013; Zhao et al., 2013), little is known about the function of the PLDγ subgroup. We therefore selected T-DNA insertion lines for the three family members *PLDγ1* (*At4g11850*), *PLDγ2* (*At4g11830*) and *PLDγ3* (*At4g11840*) (Supplemental Fig. S1A-E), which are closely related and share 93 to 95 % similarity in their amino acid sequence (Qin and Wang, 2002). These mutant lines were previously reported to have a wild type like response to infection with the barley powdery mildew fungus *Blumeria graminis* f. sp. *hordei* (Pinosa et al., 2013). To investigate whether these PLDs play a role in bacterial resistance, we infected *pldγ1-1* (Salk_066687C), *pldγ2* (Salk_078226) and *pldγ3* (Salk_084335) with the virulent bacterial strain *Pseudomonas syringae* pv*. tomato* (*Pto*) DC3000. Whereas *pldγ2* and *pldγ3* mutant plants supported bacterial growth similar to wild-type plants, genetic inactivation of *PLDγ1* resulted in significantly less growth of *Pto* DC3000 (Fig. 1A). To verify this phenotype, we infected an additional T-DNA insertion line, *pldγ1-2* (Supplemental Fig. S1A and S1C), with *Pto* DC3000. Again, in *pldγ1-2* plants bacterial growth was reduced compared to wild-type plants (Supplemental Fig. S1F), although to a lesser and more variable extent than in *pldγ1-1* plants. This could be explained by higher residual *PLDγ1* activity in *pldγ1-2* mutants compared to *pldγ1-1* (Supplemental Fig. S1C). We therefore sequenced the genome of the *pldγ1-1* mutant to determine whether this line contained secondary T-DNA insertion sites. Genome sequencing confirmed the location of the T-DNA insertion within the eighth exon of *PLDγ1* (Pinosa et al., 2013), but also revealed two additional T-DNA insertion events in the *pldγ1-1* genome, one in the promoter region of the gene *At1g77460* and the other one in the *At2g31130* promoter region, which is now also indicated on the NASC site (http://arabidopsis.info/MultiResult?name=SALK_066687&Android=F). However, mutant lines with T-DNA insertions in the coding regions of either *At1g77460* or *At2g31130* (Supplemental Fig. S2A and S2B) were similar to the wild type in growth of *Pto* DC3000 (Supplemental Fig. S2C), indicating that increased bacterial resistance of *pldγ1-1* is most likely due to genetic inactivation of *PLDγ1*. To corroborate this hypothesis further, we generated complementation lines by expressing a *PLDγ1-GFP* fusion driven by the cauliflower mosaic virus 35S promoter in the *pldγ1-1* mutant background. Independent transgenic lines with wild-type like *PLDγ1* transcript levels and PLDγ1-GFP protein expression were selected (Supplemental Fig. S3A and S3B) and T3 generation plants were infected with *Pto* DC3000. All tested transgenic *pldγ1-1/35S::PLDγ1-GFP* lines showed at least partial restoration of bacterial susceptibility (Supplemental Fig. S3C), confirming that loss of PLDγ1 is the cause for decreased bacterial growth in *pldγ1-1* mutants.

**Figure 1.**
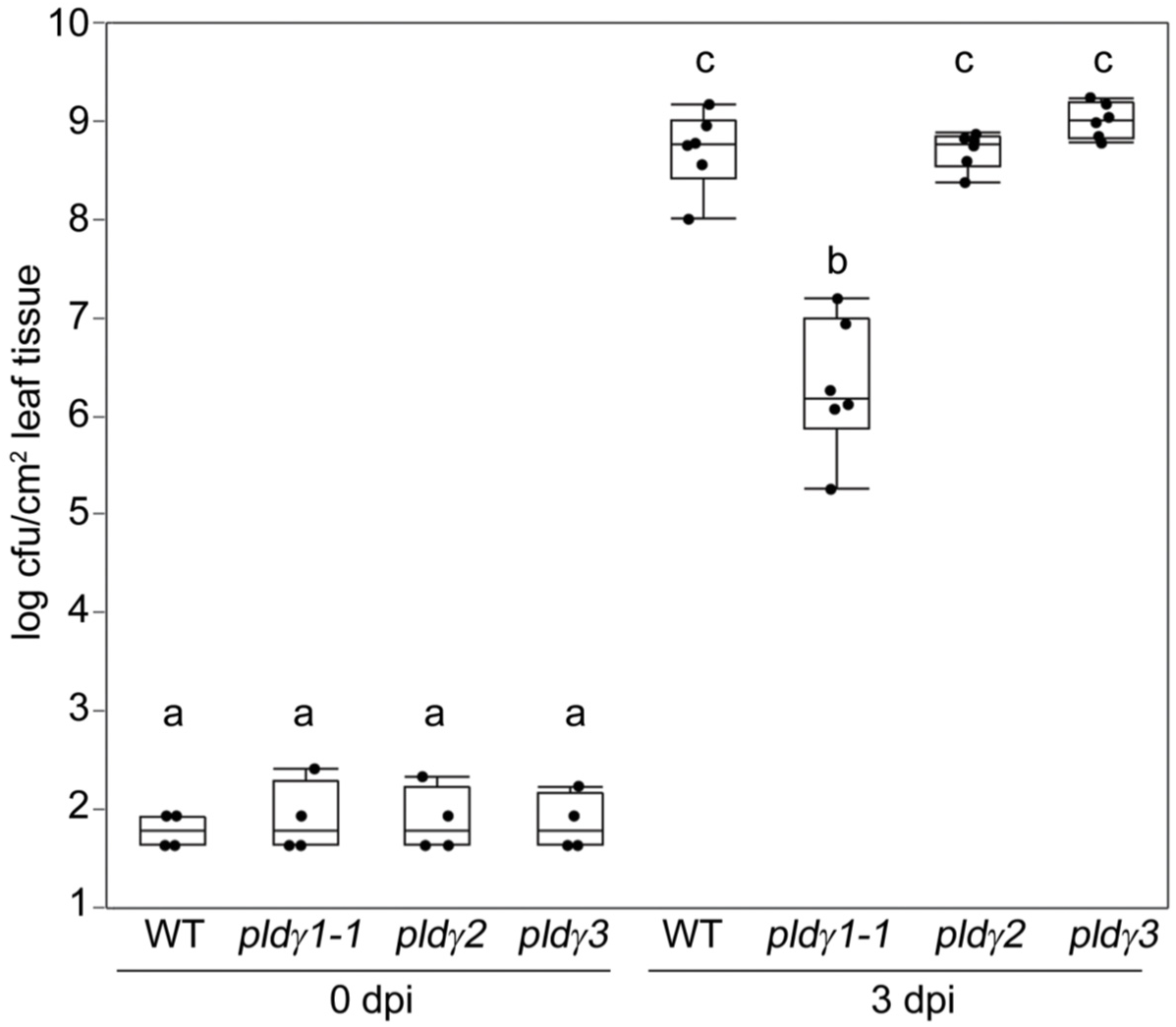
Genetic inactivation of *PLDγ1* results in an increased resistance to bacterial infection. Wild-type plants (WT) or mutant lines of *pldγ1-1*, *pldγ2* and *pldγ3* were infiltrated with 10^4^ cfu/mL of virulent *Pseudomonas syringae pv. tomato* DC3000. At day 0 and 3 post inoculation (dpi) bacterial growth was quantified by counting colony-forming units (cfu). Box plots show minimum, first quartile, median, third quartile, and a maximum of log cfu/cm^2^ leaf tissue (*n*=4 for 0 dpi; *n*=6 for 3 dpi). Labels a-c indicate homogenous groups according to post-hoc comparisons following one-way ANOVA (Tukey-Kramer multiple comparison analysis at a probability level of p < 0.05). The entire experiment was repeated three times, and the results were similar.

### Fungal resistance is enhanced in *pldγ1* mutants

We next investigated whether *pldγ1* mutants are also affected in their resistance towards the necrotrophic fungal pathogen *Botrytis cinerea*. Spore inoculation of wild-type, *pldγ2* and *pldγ3* plants resulted in comparable lesions within three days post infection and an elevated fungal DNA content in infected leaves (Supplemental Fig. S4). However, lesion sizes were much reduced in *pldγ1-1* and to a lesser extend in *pldγ1-2* mutants, which correlated with lower fungal DNA accumulation in the mutants compared to the wild type (Fig. 2 and Supplemental Fig. S4). Hence, loss of functional PLDγ1 not only resulted in increased bacterial, but also increased fungal resistance.

**Figure 2.**
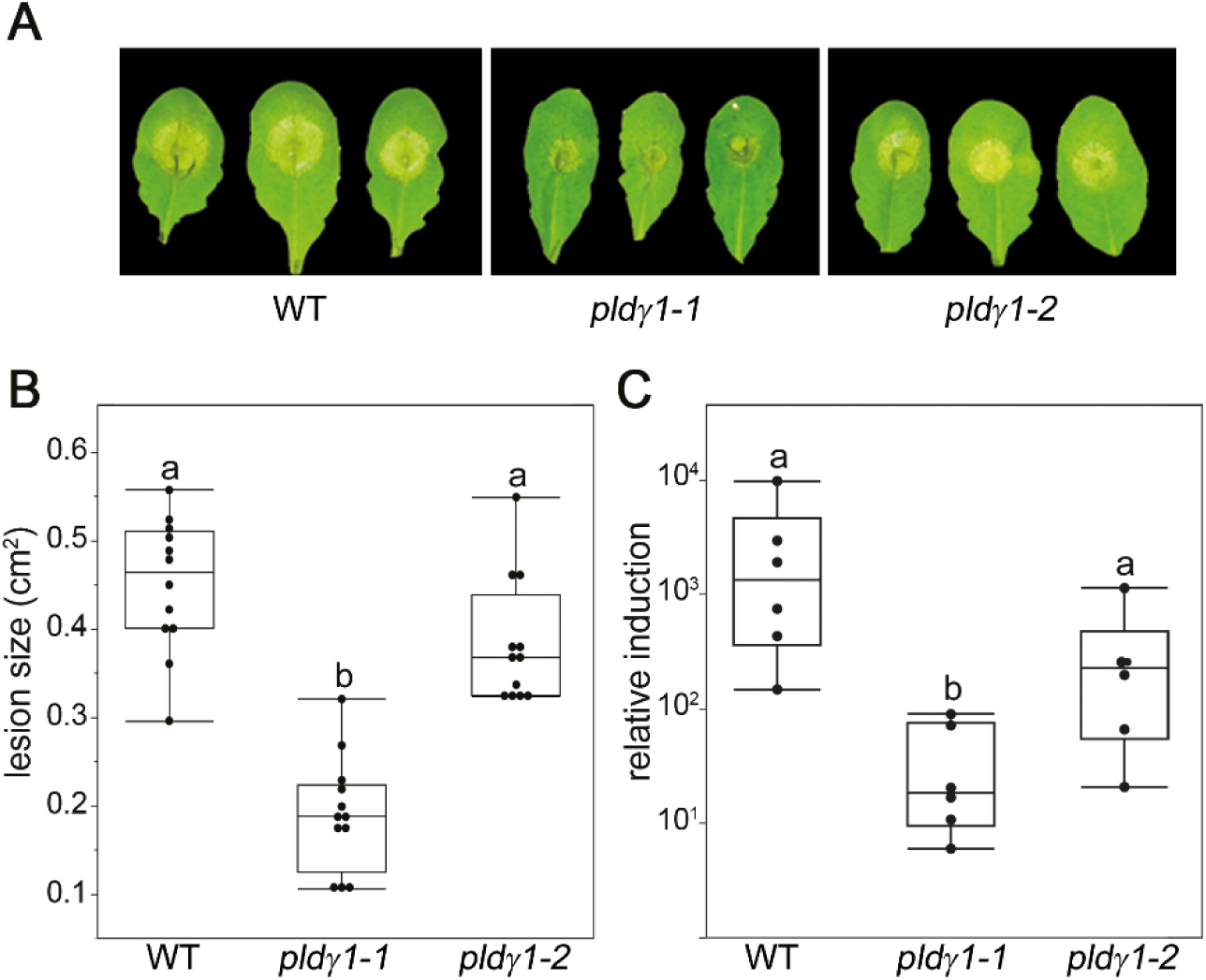
*pldγ1-1* mutants are more resistant to *Botrytis cinerea* infection. Leaves of 6-week old plants were inoculated with 5×10^6^ /mL *Botrytis cinerea* spores. A, After two days, symptom development was documented and three representative leaves per line are shown. B, Lesion sizes were determined with a pixel-based approach and then calculated using a 1 cm^2^ standard. Box plots show minimum, first quartile, median, third quartile, and a maximum (*n*=12). Labels a-b indicate homogenous groups according to post-hoc comparisons following one-way ANOVA (Tukey-Kramer multiple comparison analysis at a probability level of *p* < 0.05). B, Three days after inoculation total DNA was extracted from infected leaf material and used for the quantification of fungal biomass via RT-qPCR. The relative amount of *Botrytis cinerea* genomic *Actin-*DNA levels to Arabidopsis *Rubisco* (large subunit) levels was used to quantify fungal biomass. Box plots show minimum, first quartile, median, third quartile, and a maximum of fold induction of *Botrytis Actin* at day 3 compared to day 0 (*n*=6). Labels a-b indicate homogenous groups according to Kruskal-Wallis one-way ANOVA, followed by an each-pair comparison Wilcoxon rank-sum test with a probability level of p < 0.05. All experiments were performed three times.

### *pldγ1* mutants show an elevated oxidative burst

Increased resistance to bacterial infection has been associated with the activation of basal immunity (Zipfel et al., 2004; Zipfel et al., 2006). Hence, we tested whether *pldγ1* mutants are also altered in PTI responses. We treated mutant and wild-type plants with the MAMP flg22, a 22 amino acid long peptide of the conserved N-terminal part of flagellin that is recognized by the FLS2 immune receptor (Gómez-Gómez and Boller, 2000), and investigated the activation of mitogen-activated protein kinases (MAPKs) and the accumulation of reactive oxygen species (ROS) as two distinct branches of PTI signaling (Bigeard et al., 2015). Flg22-induced activation of particular MAPKs is an early event and believed to contribute to transcriptional reprogramming and resistance (Bethke et al., 2012). Post-translational MAPK activation was determined in Arabidopsis seedlings by Western Blot analysis using a p44/42 antibody raised against phosphorylated MAPKs. Flg22-induced activation of immunity-related MAPKs MPK6, MPK3 and MPK4/11 was indistinguishable in *pldγ1* mutants, complementation lines and wild-type plants (Figs. 3A and Supplemental Fig. S5), indicating that a loss of PLDγ1 does not impinge on the activation of this signaling cascade. Since the accumulation of ROS after flg22 stimulation is independent of MAPK activation (Zhang et al., 2007; Segonzac and Zipfel, 2011; Xu et al., 2014), we next studied the oxidative burst in our *pldγ* mutant lines. Treatment of *pldγ2* and *pldγ3* mutants, as well as the T-DNA insertion lines for *At1g77460* and *At2g31130*, with flg22 resulted in ROS responses similar to those observed in wild-type plants (Supplemental Figs. S2D and S6A). However, ROS levels in *pldγ1-1* were significantly increased, and reached about double the amount of wild-type, *pldγ2-* and *pldγ3* plants, as well as *pldγ1/35S::PLDγ1-GFP* complementation lines (Fig. 3B, Supplemental Fig. S6). Together with the increased bacterial resistance only observed in *pldγ1* mutants, these results suggest that PLDγ1 acts as a negative regulator of plant immunity.

**Figure 3.**
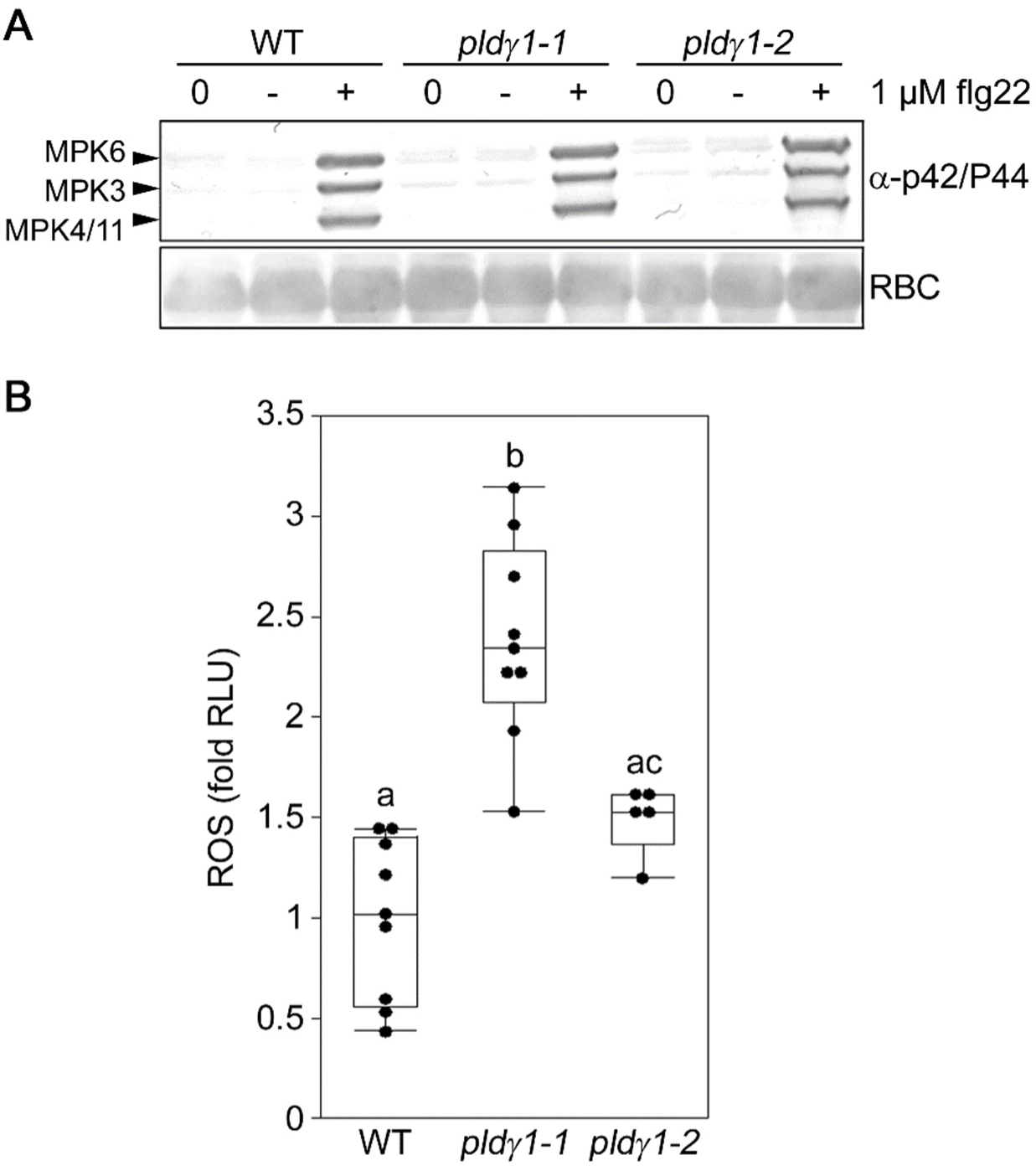
ROS levels, but not MAPK activation, are affected in *pldγ1-1* mutants. A, 10-day old seedlings of the wild type or the *pldγ1-1* and *pldγ1-2* mutants were collected at time point zero (0) or were treated for 15 min with 1 μM flg22 (+) or water as control (-). The activation of MAPKs was visualized by Western Blot analysis using the phospho-p44/42 MAP kinase antibody. Ponceau S Red-staining of the membrane served as a loading control (RBC, Ribulose-bis-phosphate-carboxylase large subunit). B, Arabidopsis leaf pieces of the indicated mutant line were treated with 1 µM flg22 or water as control, and ROS production was monitored over time. Shown are relative light units (RLU) expressed as fold induction to the mean of the wild type, which was set as 1. Box plots show the minimum, first quartile, median, third quartile, and a maximum of fold induction of peak value minus background value (*n*≥6). Water-treated samples had no detectable ROS production and are therefore not displayed in the figure. Labels a-c indicate homogenous groups according to post-hoc comparisons following one-way ANOVA (Tukey-Kramer multiple comparison analysis at a probability level of p < 0.05). All experiments were performed at least three times.

### *pldγ1* mutation does not affect phytohormone levels

Plant hormones play an important role in immunity (Robert-Seilaniantz et al., 2011; Pieterse et al., 2012; Ku et al., 2018), especially salicylic acid (SA) and jasmonic acid (JA), which are biosynthetically induced after pathogen attack, resulting in activation of downstream pathways (Robert-Seilaniantz et al., 2011; Pieterse et al., 2012). SA is often correlated with biotrophic infection while JA is typically associated with resistance towards necrotrophic pathogens (Pieterse et al., 2012). We therefore determined SA- and JA-levels in untreated *pldγ1-1* mutant plants. Neither basal SA- and JA-levels in *pldγ1-1* lines were distinguishable from those obtained from wild-type plants (Supplemental Fig. S7A and B). Likewise, gene expression levels of SA- and JA-marker genes after treatment with flg22 were not affected by genetic inactivation of *PLDγ1* (Supplemental Fig. S7C). Hence, PLDγ1 is unlikely to exert its function in immunity via SA- or JA-signaling pathways.

### PA levels are unaltered in *pldγ1* mutants

PLDs can cleave membrane phospholipids to produce phosphatidic acid (PA) as lipid second messenger (Testerink and Munnik, 2011). Hence, we analysed basal PA levels in our *pldγ1* mutants. To this end, phospholipids in Arabidopsis seedlings were metabolically labelled with ^32^P-P_i_, and ^32^P-PA levels were determined after extraction and thin layer chromatography. No difference between wild-type and *pldγ1* seedlings in basal PA levels were found that would explain their difference in bacterial resistance (Fig. 4). Upon flg22 treatment, a small increase in PA was observed but this was the same for both wild-type and *pldγ1* plants (Fig. 4A). Salt stress resulted in a stronger PA accumulation, reaching about 1 % of total phospholipids (Bargmann et al., 2009; Yu et al., 2010), but also there no difference between wild type and mutant was observed (Fig. 4B).These results indicate that, at least in our experimental set up, the genetic inactivation of *PLDγ1* does not impinge on PA levels, which are therefore not likely to be the cause for the observed increase in bacterial resistance of the *pldγ1* mutant.

**Figure 4.**
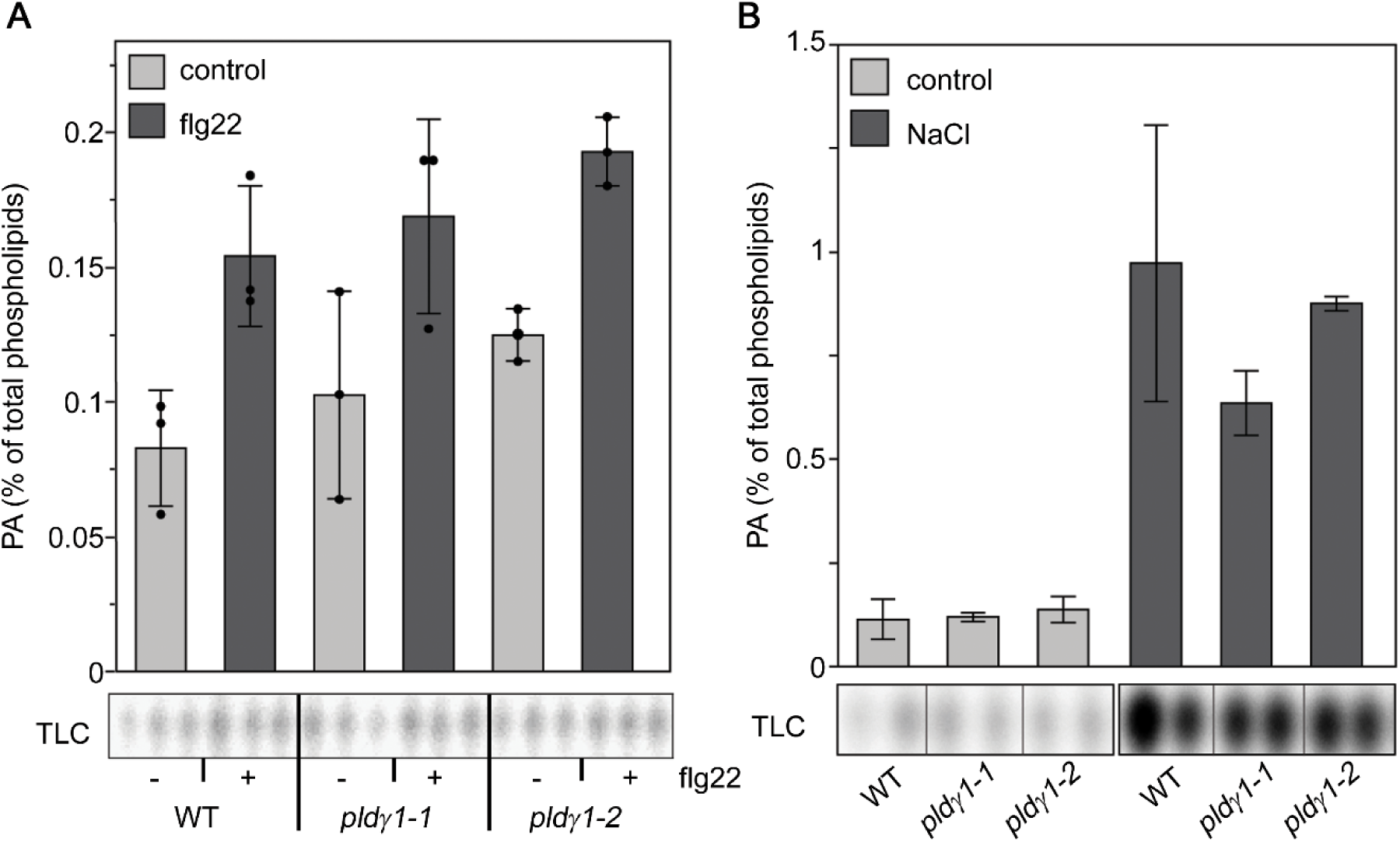
PA levels are not affected by PLDγ1 depletion. 5-day old seedlings of wild type, *pldγ1-1* or *pldγ1-2* were labelled with ^32^P_i_ for 16 h and then treated for 15 minutes with either 1 µM flg22 (A) or 300 mM NaCl (B) or cell-free medium (control) as indicated. Lipids were extracted, separated by EtAc-TLC and the radioactivity incorporated into PA was quantified by phosphoimaging (bottom panels). Data (top panels) represent the average of 2-3 biological replicates and are expressed in relation to the radioactivity of the total phospholipids.

### PLDγ1 associates with BIR2 and BIR3

Since PA levels appeared to be unaffected by genetic inactivation of *PLDγ1*, we searched for an alternative molecular mechanism as to how PLDγ1 negatively regulates the plant immune response. Intriguingly, the *pldγ1* phenotype (i.e. elevated resistance to bacteria and increased production of ROS upon flg22 treatment, Figs 1, 3B and Supplemental Fig. S6C) is reminiscent of the behaviour of *bir2* mutants (Halter et al., 2014). The BAK1 interacting-receptor kinase BIR2 and its close homolog BIR3 act as negative regulators of plant immune responses by constitutively interacting with the co-receptor BAK1, thereby preventing its interaction with ligand-binding LRR-RLKs, such as FLS2 or the brassinolide-receptor BRI1 (Halter et al., 2014; Imkampe et al., 2017).

Therefore, we first investigated whether the subcellular protein localization would permit PLDγ1 to associate with BIR2/3 proteins at the plasma membrane (PM). As the fluorescence signal in the *pldγ1-1/35S::PLDγ1-GFP* complementation line was too weak for microscopic detection, we transiently expressed PLDγ1-GFP together with either BIR2-RFP or BIR3-CFP in *Nicotiana benthamiana*. Using confocal microscopy, we could clearly detect GFP, RFP and CFP signals localized to the PM of leaf epidermal cells (Fig. 5), confirming that PLDγ is localised at the PM, most likely due to N-terminal myristoylation (Qin et al., 1997; Fan et al., 1999). Moreover, an overlay of fluorescent signals indicated co-localization of PLDγ1 with BIR2 and BIR3, respectively, at the PM in tobacco plants.

**Figure 5.**
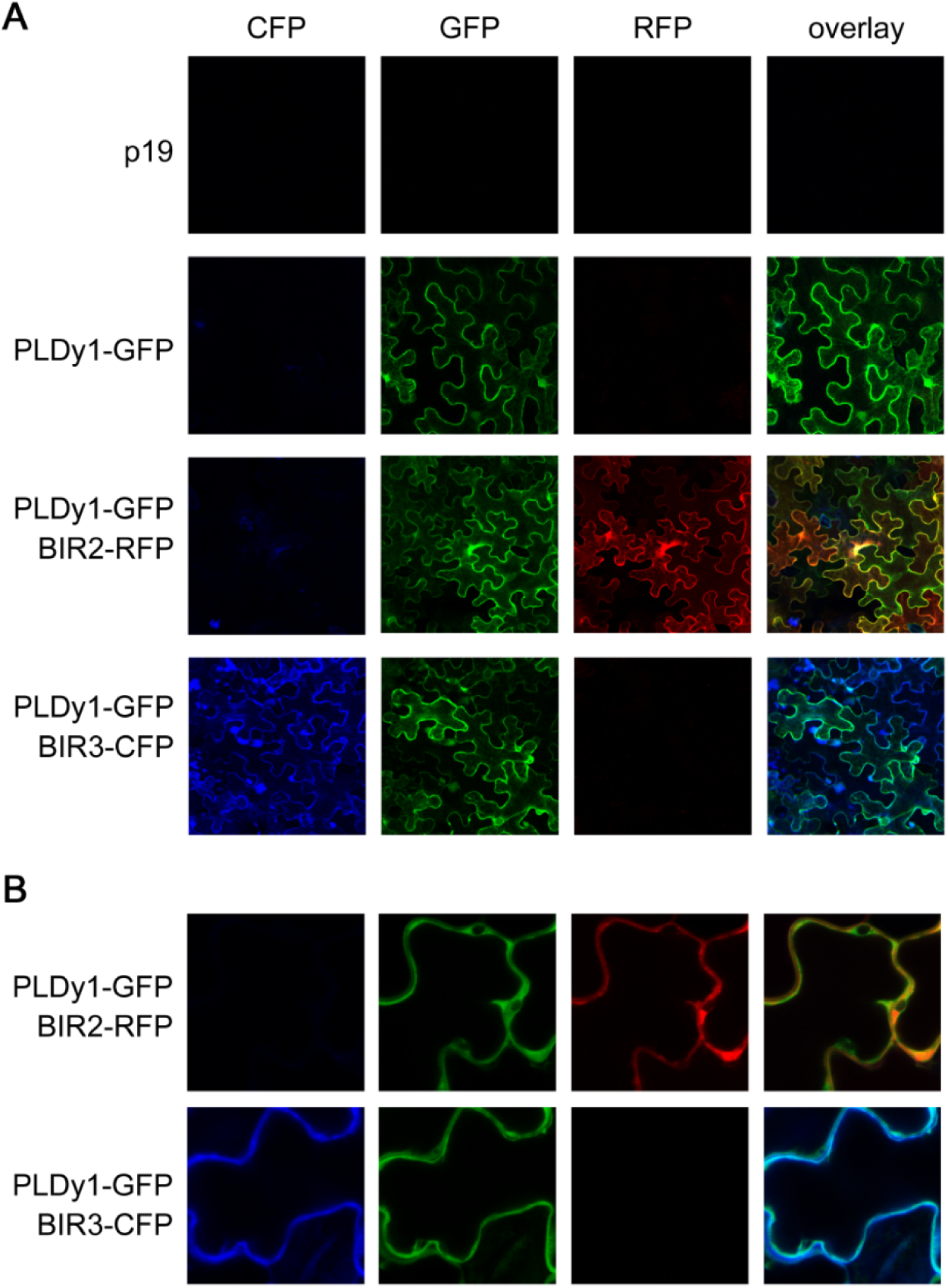
PLDγ1 co-localizes with BIR2 and BIR3 at the plasma membrane. A, PLDγ1 was transiently expressed in *Nicotiana benthamiana* as GFP-fusion either alone or together with BIR2-RFP or BIR3-CFP. Fluorescence in epidermal cells was monitored at day 3 using confocal laser-scanning microscopy. B, Close-up view of the samples shown in (A).

To test whether PLDγ1 could interact with BIR2 or BIR3, we transiently expressed PLDγ1 fused to a C-terminal GFP-tag together with either BIR2 or BIR3 fused to a C-terminal myc-epitope in *N. benthamiana* leaves. Co-immunoprecipitation with protein extracts from those leaves revealed that PLDγ1 can be found in a constitutive complex with both BIR2 and BIR3 (Fig. 6), regardless of which protein partner was pulled down with the corresponding affinity beads. Interestingly, treatment with flg22 resulted in more BIR2-myc co-purified with PLDγ1-GFP compared to the control treatment, whereas the PLDγ1-BIR3 interaction seemed to be largely unaffected by elicitation (Fig. 6). Quantification of immunoprecipitated BIR2 protein levels relative to PLDγ1 signals showed that within 5 min of flg22 treatment about 8.65 times more BIR2-myc could be found in complex with PLDγ1-GFP (when myc-beads were used for immunoprecipitation) and, likewise, 2.42 times more PLDγ1-GFP could be found in complex with BIR2-myc (when GFP-beads were used for immunoprecipitation), indicating a ligand-dependent enhancement of the PLDγ1/BIR2 protein interaction. PLDy1 complex formation with BIR2 and BIR3 could also be confirmed in experiments exchanging epitope tags, and again more PLDγ1 could be found in BIR2 precipitates in the presence of flg22 (Fig. 6B, C, E and F). Association of PLDγ1 with BIR2 and BIR3 was specific, as PLDγ1 did not co-purify with other LRR-RLKs such as FLS2 or BAK1 (Supplemental Fig. S8).

**Figure 6.**
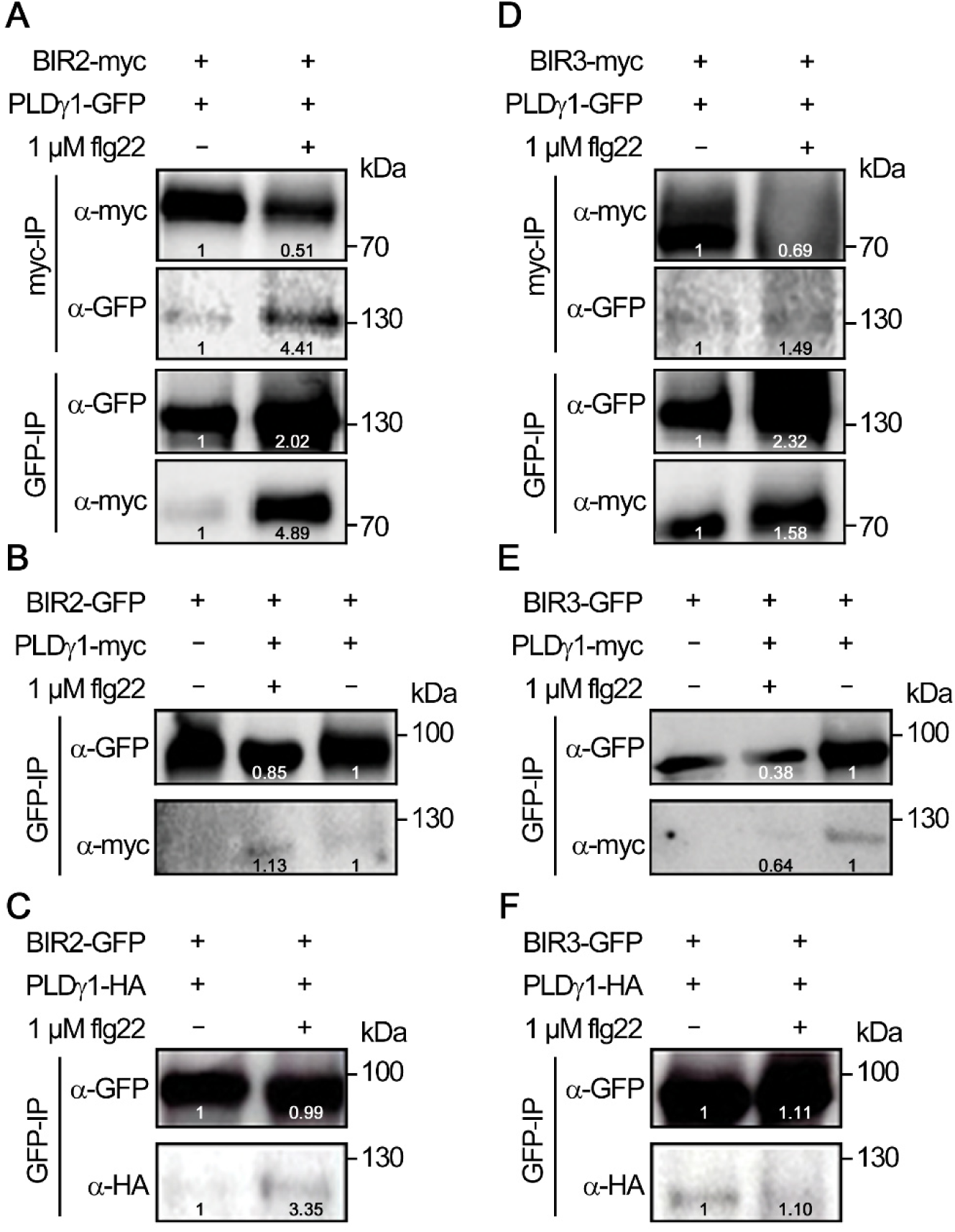
PLDγ1 can be found in complex with BIR2 and BIR3. Western Blot analysis of transiently expressed proteins in *N. benthamiana* three days after infiltration. Leaf material was harvested 5 minutes after treatment with 1 µM flg22 (+) or water (-). After protein extraction the proteins were subjected to immunoprecipitation with GFP- or myc-affinity beads as indicated. For different immunoprecipitations within one experiment (A and D) the same source material was used. Immunoprecipitated and co-purified proteins were detected with tag-specific antibodies as indicated. A and D, Co-immunoprecipitation of BIR2-myc (A) or BIR3-myc (D) and PLDγ1-GFP using myc-trap beads or GFP-trap beads, respectively. B and E, Co-purification of BIR2-GFP (B) or BIR3-GFP (E) and PLDγ1-myc precipitated with GFP-trap beads. C and F, Co-purification of BIR2-GFP (C) or BIR3-GFP (F) and PLDγ1-HA precipitated with GFP-trap beads. Inserted numbers indicate quantification of signal intensities. All experiments were repeated at least three times with similar results.

To investigate whether PLDγ1 could directly physically interact with BIR2 and BIR3 we performed bimolecular fluorescence complementation (BiFC) analysis. Although this system confirmed interaction of the Arabidopsis Calmodulin AtCaM1 with the *Phytophthora infestans* RXLR effector SFI5 (Zheng et al., 2018), we could, however, not detect any direct interaction between PLDγ1 and the BIR2 and BIR3 full-length proteins *in planta* (Supplemental Fig S9). This result indicates that although PLDγ1 is present in the same protein complex as BIR2 and BIR3, these proteins most likely do not come into direct contact with each other.

## Discussion

PLDs are lipid-hydrolyzing enzymes with multiple functions during plant abiotic and biotic stress responses as well as cellular and developmental processes (Bargmann and Munnik, 2006; Zhao, 2015; Hong et al., 2016; Takáč et al., 2019). A direct product of the hydrolytic activity of PLDs towards plasma membrane (PM) lipids is PA, a typical lipid second messenger which can bind to and modulate the activity of proteins such as kinases and phosphatases, 14-3-3 proteins, transcription factors or proteins involved in calcium signaling, oxidative burst, endocytosis or cytoskeleton assembly (reviewed in (Munnik, 2001; Wang, 2004; Zhao, 2015; Pokotylo et al., 2018; Takáč et al., 2019)). Notably, PA-binding to a protein can not only affect its enzymatic activity but also its localization where PA-binding mediates membrane-tethering of soluble proteins (Pokotylo et al., 2018).

### PA could act as both negative and positive regulator of plant immunity

Upon pathogen-infection or elicitor treatment, PA rapidly accumulates in leaf tissues and suspension cells (van der Luit et al., 2000; Laxalt et al., 2001; den Hartog et al., 2003; de Jong et al., 2004; Bargmann et al., 2006; Zhao et al., 2013) and is assumed to be an early signaling molecule with generally positive regulatory function. For instance, exogenously applied PA can induce ROS production, MAPK and defence gene activation, and ultimately cell death (de Jong et al., 2004; Park et al., 2004; Andersson et al., 2006; Yu et al., 2010). However, genetic inactivation of PLD isoforms such as *PLDβ1* (Bargmann et al., 2006; Yamaguchi et al., 2009; Zhao et al., 2013) and *PLDγ1* (Figs 1 and 2) resulted in increased immunity, suggesting that PA can also act as negative regulator in plant defence. Notably, fatty acid compositions of PLD-produced PA vary depending on the localization of the respective PLD isoform and PA-protein interaction might be determined by the availability of the corresponding target protein in a given tissue at a given developmental stage or environmental condition (Testerink and Munnik, 2005; Pokotylo et al., 2018), possibly enabling PA to exert both positive and negative regulatory roles. In addition to PLD, PA can also be produced via the combined action of PLC and DGK (Arisz et al., 2009). At least in cell suspensions, this is the major pathway for flg22-triggered PA production (van der Luit et al., 2000), supporting our observation that the slight flg22-induced PA accumulation was not affected by the *pldγ1* mutation (Fig. 4). Nevertheless, PLDs might have a more significant role in PA production after stimulation with other MAMPs such as fungal xylanase (van der Luit et al., 2000) or in other pathways such as effector triggered-immunity (de Jong et al., 2004; Andersson et al., 2006). In any case, a direct negative function of PA in immunity remains to be demonstrated.

### PLDs might have a function in receptor localization and stability

Apart from producing PA as signaling molecule, phospholipases are also involved in remodeling membrane lipids, which affects the biophysical properties and hence membrane functionality (Platre et al., 2018; Mamode Cassim et al., 2019). The PM is organized in highly dynamic sub-compartments, so called micro- or nanodomains, depending on size and shape, which allow the accommodation of specific protein clusters (Malinsky et al., 2013; Ott, 2017). Protein clustering within the PM has been demonstrated for the immune receptor FLS2 (Bücherl et al., 2017; McKenna et al., 2019) as well as its co-receptor BAK1 (Hutten et al., 2017). Importantly, the stability of FLS2-protein clusters depends on the lipid composition of the surrounding membrane patch (Bücherl et al., 2017). Likewise, the mammalian receptor tyrosine kinase EPIDERMAL GROWTH FACTOR RECEPTOR (EGFR), is found in PM nanodomains, and formation of these protein clusters is dependent on the integrity of PM lipid composition (Gao et al., 2015). Moreover, both FLS2 and EGFR are rapidly internalized upon ligand-binding (Haigler et al., 1978; Robatzek et al., 2006). EGFR endocytosis and subsequent degradation depends on PLD activity (Shen et al., 2001), linking PLD action with receptor internalization (Donaldson, 2009). The diffusion rate of FLS2 within the PM as well as FLS2 endocytosis is dependent on an intact cytoskleleton assembly (Robatzek et al., 2006; McKenna et al., 2019). Cytoskleletal dynamics in turn are altered by PLDs, such as PLDβ1 and PLDδ, which bind to actin and microtubules and PLD enzymatic activity is regulated by this protein-protein interaction (Munnik and Musgrave, 2001; Dhonukshe et al., 2003; Kusner et al., 2003; Pleskot et al., 2010; Zhao, 2015). Hence, PLD-mediated membrane lipid remodelling or PLD-cytoskeleton interactions might impinge on FLS2 (or general immune receptor) protein distribution or stability at the PM or receptor internalization upon activation.

### PLDγ1 is a novel complex partner for the negative regulator BIR2

PA rapidly accumulates in pathogen-infected plants, apparently to regulate immune responses via PA-protein interactions. In addition, PLD proteins themselves can directly interact with a range of proteins, indicating additional PLD-functions apart from lipid hydrolysis. One example are the aforementioned cytoskeleton components. PLDα1, as another example, interacts with the heterotrimeric G protein GPA1 and promotes the exchange of GTP to GDP, thereby mediating ABA-induced inhibition of stomatal opening (Mishra et al., 2006). PLDα1 also complexes with the cysteine-rich receptor-like kinase 2 (CRK2) and affects its relocation within the PM upon salt stress (Hunter et al., 2019). Intriguingly, PLDα1 was also found to physically interact with yet another protein kinase, MPK3, and *pldα1 mpk3* double mutants displayed an increased tolerance to salinity and ABA treatment compared to the single mutants (Vadovič et al., 2019), indicating multiple sites of action for PLDα1 in Arabidopsis salt stress and ABA signaling. In plant immunity, PLDδ physically interacts with the immune receptor *Resistance to Pseudomonas syringae pv. maculicola 1* (RPM1) and negatively regulates its function (Yuan et al., 2019).

We found PLDγ1 in complex with the LRR-RLKs BIR2 and BIR3, which function as negative regulators of immune receptors (Halter et al., 2014; Imkampe et al., 2017). Interestingly, we observed differences in PLDγ1 complex formation: whereas flg22 treatment resulted in further BIR2 recruitment, the PLDγ1/BIR3 complex remained largely unaffected upon elicitation (Fig. 6). Notably, although both BIR2 and BIR3 interact with BAK1 to hinder this co-receptor from joining receptor complexes without stimulus, several other behavioral differences for BIR2 and BIR3 have been reported (Halter et al., 2014; Imkampe et al., 2017): i) BIR3, but not BIR2, directly interacts with ligand-binding receptor kinases such as FLS2 or BRI1; ii) BIR3, but not BIR2, is a negative regulator of BRI1-mediated brassinosteroid-perception, iii) *bir2*, but not *bir3* mutants, show increased resistance towards bacterial infection, iv) BIR2 is the better phosphorylation target of BAK1. This suggests that BIR3, although having several functions in common with BIR2, has additional distinct functions and might act at least partially via a different molecular mechanism than BIR2. Importantly, the *pldγ1* mutant phenotype with respect to bacterial resistance mirrored the behavior of *bir2*, but not *bir3* mutants, making BIR2 the more likely complex partner for PLDγ1. As we could not observe any association of PLDγ1 with BAK1 or FLS2 (Supplemental Fig. S8), and as PLDγ1/BIR2 interaction was enhanced by flg22 (Fig. 6), PLDγ1 is likely recruited to BIR2 after its release from BAK1. Future work will now be needed to elucidate the exact mechanism and function of PLDγ1 complex formation with BIR2 and BIR3.

### Conclusions

PLDγ1 is a novel negative regulator of plant immune responses. We hypothesize that PLDs shape the lipid composition of membranes to allow the formation of distinct PM nanodomains to aid the relocation of immune-receptors and/or complex formation with their positive and negative regulators (such as BIR2 and BIR3). Changes in lipid composition or PLD hydrolytic activity might additionally influence the association of cytoplasmic proteins with the PM or the dissociation of membrane-tethered proteins from the PM upon signaling (e.g. RLCKs). PLD actions on PM lipids are most likely very local (within micro- or nanodomains) and thus do not necessarily result in measurable alterations in PA levels (we did not detect any changes in PA amounts in *pldγ1* mutants). Hence, in addition to being merely producers of PA, PLDs including PLDy1 might have novel functions in regulating cellular responses via cluster formation with immunity-related proteins.

## MATERIALS AND METHODS

### Plant Material

*Arabidopsis thaliana* plants were grown on soil or half-strength Murashige and Skoog (MS) medium as described (Brock et al., 2010). Plants were grown in climate chambers under short-day conditions (8 h light/16 h darkness, 150 μmol/cm^2^s white fluorescent light, 40-60 % humidity, 22 °C). Arabidopsis accession Col-0 was the background for all mutants used in this study, genotyping was performed with primers listed in Supplemental Table S1. Mutant lines *pldγ1-1* (Salk_066687C), *pldγ2* (Salk_078226), *pldγ3* (Salk_084335) and *bir2-1* (GK-793F12) have been previously described (Pinosa et al., 2013; Halter et al., 2014) and are listed in Supplemental Table S1, together with *pldγ1-2* (GK-264A03) and T-DNA insertion lines selected for *At1g77460* and *At2g31130*. *Nicotiana benthamiana* plants were grown in the greenhouse (16 h light, 22 °C).

### Generation of Constructs and Transgenic Plants

For co-immunoprecipitation experiments, all C-terminal epitope-tag fusion constructs were made by cloning full length coding sequences as PCR fragments first into pCR8/GW/TOPO and afterwards into vectors pGWB5 (GFP-tag), pGWB14 (3xHA) or pGWB17 (4xmyc) (Nakagawa et al., 2007) using the Gateway® LR Clonase® II Enzyme Mix (Thermo Scientific). For co-localization, full length coding sequences were recombined into vectors pB7RWG2.0 (BIR2-RFP), pH7CWG2.0 (BIR3-CFP) or pGWB5 (PLDγ1-GFP) (Karimi et al., 2005; Nakagawa et al., 2007).

For BiFC analyses, full length coding sequences of the respective genes were cloned into pDONR221P1P4 and pDONR221P3P2 (Thermo Scientific) followed by recombination into the pBiFCt-2in1-CC vector (Grefen and Blatt, 2012) to generate C-terminal fusions with the C- and N-terminal part of YFP, respectively.

Complementation lines were generated by the *Agrobacterium*-mediated floral dip method (Clough and Bent, 1998) of *pldγ1-1* mutant lines using the full length coding sequence of *PLDγ1* fused to a C-terminal GFP-tag in the binary vector pGWB5 (Nakagawa et al., 2007). Restored gene transcription in transformed seedlings was verified by RT-qPCR and Western Blot analyses, and plants were used for infection experiments in the T3 generation.

### Plant Infections

*Pseudomonas syringae* pv. *tomato* DC3000 (*Pto* DC3000) was grown overnight in King’s B medium, centrifuged, washed and diluted in 10 mM MgCl_2_ to a density of 10^4^ cfu mL^−1^. Bacteria were pressure-infiltrated into leaves of 5 to 6-week old Arabidopsis plants. Leaves were harvested at day 0 and 3, surface sterilized in 70 % (v/v) ethanol and washed in ddH_2_0 for 1 min each. Two leaf discs (5 mm diameter) per leaf were ground in 200 μL of a 10 mM MgCl_2_ solution, diluted serially and plated on LB plates containing rifampicin and cycloheximide. After 2 days of incubation at 28°C colony-forming units were counted.

Spores of *Botrytis cinerea* isolate B0-10 were diluted to a final concentration of 5×10^6^ spores mL^−1^ in PDB medium and a 5 µL drop per leaf was used to inoculate 5 to 6-week old Arabidopsis plants. Lesion sizes were determined after 2 days using the Photoshop CS5 Lasso tool. Selected pixels were counted and the lesion size in cm^2^ was calculated using a 0.5 cm^2^ standard. Fungal biomass quantification was performed by extracting DNA from infected leaf material. 3 infected leaves/plant were frozen in liquid nitrogen using screw cap vials containing a ceramic bead mix (2 mm and 0.5 mm diameter). Tissue was homogenized using the Precellys 24 tissue homogenizer (Bertin Instruments) for 2 × 30 sec at room temperature. After addition of 300 µL extraction buffer (2.5 mM LiCl, 50 mM Tris-HCl pH 8, 62.5 mM EDTA 4 % (v/v) Triton X100), samples were incubated for 10 min at room temperature, followed by purification using phenol-chloroform-extraction. Genomic DNA was finally dissolved in nuclease-free water and subjected to RT-qPCR using primers listed in Supplemental Table S2.

### Detection of immune responses

The detection of reactive oxygen species (ROS) in leaf pieces of 5 to 6-week old Arabidopsis plants was measured using 96-well plates containing in a final volume of 100 µL two leaf pieces, 5 µg mL^−1^ horseradish peroxidase (Applichem) and 20 µM luminol L-012 (Wako Chemicals, USA) and the elicitor to be tested per well as described (Albert et al., 2010). Light emission was measured as relative light units in a 96-well luminometer (Mithras LB 940; Berthold Technologies). Immunoblot analyses using the anti-phospho p44/42 (Erk1/2) MAP kinase antibody (Cell signaling technology) was done with 10-day old seedlings as described (Brock et al., 2010).

### RNA extraction and reverse transcription-quantitative PCR (RT-qPCR)

Total RNA from 100 mg leave tissue (5 to 6-week old Arabidopsis plants) or 10 nine-day old seedlings was purified using the NucleoSpin® RNA Plus kit (Macherey-Nagel). First-strand cDNA synthesis was performed from 1 µg of total RNA using RevertAidTM MuLV reverse transcriptase (Thermo Scientific). Quantitative PCR reactions and measurements were performed with the iQ5 Multi-color real-time PCR detection system (Bio-Rad) using the Maxima SYBR Green/Fluorescein qPCR Master Mix (Thermo Scientific) and gene specific primers listed in Supplemental Table S2. Relative gene expression was calculated according to the 2^−Δct^ method (Livak and Schmittgen, 2001) to the housekeeping gene *EF-1α*.

### Determination of phosphatidic acid, salicylic acid and jasmonic acid

Analysis of phospholipids was essentially done as previously described (Munnik and Zarza, 2013). In brief, 5-day old seedlings were labelled overnight with ^32^P-orthophosphate and treated the next day with buffer with or without flg22 or NaCl. Lipids were subsequently extracted and analysed by thin layer chromatography using an ethyl acetate solvent system, composed of the organic upper phase of an ethyl acetate/iso-octane/HAc/H_2_O mixture of 13:2:3:10 (v/v) (Munnik and Laxalt, 2013). Radio-labelled phospholipids were visualized and quantified using a PhosphoImager (GE Healthcare) and the program QuantityOne (Biorad).

For salicylic acid (SA) and jasmonic acid (JA) quantification, 200 mg leaf material of 8-week old Arabidopsis plants was frozen. The amount of SA and JA were measured by gas chromatography coupled to mass spectrometry as previously described (Wan et al., 2018), but using a. Shimadzu TQ8040 triple quad mass spectrometer (Shimadzu Cooperation), with splitless injection mode and a SH-Rxi-17SIL-MS column (30 m, 0.25 mm internal diameter, 0.25 µm film, Shimadzu Cooperation). The GC oven temperature was held initially at 70°C, then ramped at 25°C min^−1^ to 280°C and held for 7 min, then ramped at 10°C min^−1^ to 300°C and afterwards held for an additional 3 min at 300°C. Helium was used as carrier gas with a flow rate of 0.86 ml min^−1^. The mass spectrometer was operated in electron impact ionization (EI) and multiple reaction monitoring (MRM) mode.

### Protein Interaction studies

For transient expression of epitope-tagged proteins, plasmid constructs were inserted into *Agrobacterium tumefaciens* strain GV3101 and overnight cultures were used for leaf infiltration of *N. benthamiana* at O.D._600_ of 0.5 as described (Brock et al., 2010). Leaf material was harvested after three days. For protein extraction 200 mg ground tissue was resuspended in 1.6 mL solubilization buffer (25 mM Tris-HCl pH 8.0, 150 mM NaCl, 1 % (v/v) NP40, 0.5 % (w/v) DOC, 2 mM DTT and 1 tablet of "cOmplete ULTRA Tablets, Mini, EASYpack" (Roche) per 10 mL). Proteins were solubilized for 1h at 4°C with head-over-head shaking and the cell debris was removed by two times centrifugation at 20,000 g for 10 min at 4°C. For immunoprecipitation, each supernatant was incubated with pre-washed and in solubilization buffer equilibrated affinity-beads (GFP-trap or myc-trap, ChromoTek) for 1 h at 4°C with head-over-head rotation at 6 rpm. Beads were washed twice with solubilization buffer and twice with washing buffer (25 mM Tris-HCl pH 8.0, 150 mM NaCl, 2 mM DTT) before adding SDS-PAGE loading dye. Samples were analyzed by Western Blotting using the following antibody dilutions: anti-GFP (1:10,000, from goat, Sicgen), anti-myc (1:5,000, from rabbit, Sigma), anti-HA (1:5,000, from mouse, Sigma), anti-rabbit lgG-HRP-conjugate (1:10,000, from goat, Sigma), anti-goat lgG-HRP-conjugate and anti-mouse lgG-HRP-conjugate (each 1:10,000, from rabbit, Sigma).

For BiFC analyses, proteins fused to the C- and N-terminal part of YFP were transiently expressed in *N. benthamiana* and fluorescence of reconstituted YFP was visualized by confocal laser-scanning microscopy 3 days after *Agrobacterium*-infiltration.

### Whole genome sequencing

Genomic DNA of the *pldγ1-1* was isolated using the DNeasy Plant Kit (Qiagen, Germany). The library was prepared using a modified Nextera protocol (Karasov et al., 2018) and was sequenced using a HiSeq3000 Illumina^□^ platform (Illumina^□^, USA). Forty-fold genome coverage was obtained. The raw reads were trimmed using the SKEWER (v0.2.2) software (minimum quality 20 and minimum length 30) (Jiang et al., 2014) and the processed reads were mapped against the expected pROK2 insertion using the BWA MEM (v0.7.15) algorithm (Li and Durbin, 2009). Downstream processing of the files was performed using SAMTOOLS (v1.3.1) (Li et al., 2009) and PCR duplicates were removed using the PICARD (v2.2.1) algorithm (http://broadinstitute.github.io/picard). SAMTOOLS (v1.3.1) was used for sub-setting reads that were mapped at the first or last 100 nucleotides of the pROK2 insertion (Li et al., 2009). These reads were mapped to the *A. thaliana* TAIR10 reference using the BWA MEM (v0.7.15) algorithm (Li and Durbin, 2009). As previously, the newly generated file was processed using SAMTOOLS (v1.3.1) (Li et al., 2009). The file was evaluated for multiple integration points of the transgene and visualised on IGV (Thorvaldsdóttir et al., 2013).

### Statistical analysis

To test for significant differences in different experiments first the distribution of the data was determined. For parametric data one-way ANOVA followed by Tukey-Kramer multiple comparison analysis at a probability level of p < 0.05 was applied. For non-parametric data Kruskal-Wallis one-way ANOVA, followed by an each-pair comparison Wilcoxon rank-sum test with a probability level of p < 0.05 was used. All data analyses were performed by using JMP 14 software.

### Accession Numbers

Sequence data from this article can be found in TAIR under the following accession numbers: *PLDγ1* (At4g11850), *PLDγ2* (At4g11830), *PLDγ3* (At4g11840), *BIR2* (At3g28450), *BIR3* (At1g27190), *AtCaM1* (A5g37780). *SFI5* (PITG_13628). Accession numbers for all other genes/proteins used in this work are indicated in Supplemental Tables S1 and S2.

## Supplemental Material

The following supplemental materials are available.

**Supplemental Figure S1.** Characterization of T-DNA insertion lines of PLDγ family members.

**Supplemental Figure S2.** Characterization of secondary mutations in the *pldγ1-1* mutant.

**Supplemental Figure S3.** Characterization of *pldγ1-1* complementation lines.

**Supplemental Figure S4.** *pldγ1* mutants of are more resistant to *Botrytis cinerea* infection compared to *pldγ2* and *pldγ3* mutants.

**Supplemental Figure S5.** Flg22-induced MAP kinase activation is not altered in *pldγ1* mutants or complementation lines.

**Supplemental Figure S6**. ROS levels in *pldγ1-1* and *bir2* mutants are elevated compared to *pldγ2* and *pldγ3* mutant lines or *pldγ1-1* complementation lines.

**Supplemental Figure S7.** PLDγ1 depletion does not affect SA and JA-signaling.

**Supplemental Figure S8.** PLDγ1 does not interact with FLS2 and BAK1.

**Supplemental Figure S9.** PLDγ1 does not directly interact with BIR2 and BIR3 in BiFC.

**Supplemental Table S1.** Primers used for genotyping T-DNA insertion lines.

**Supplemental Table S2.** Primers used in qPCR analyses.

## ACKNOWLEDGMENTS

We are grateful to Raffaele Del Corvo for his excellent support on the characterization of secondary insertion sites, Frédéric Brunner and Xiangzi Zheng for providing the BiFC constructs for AtCaM1 and SFI5, all students of the practical course ‘Modern Genetic Engineering 2018’ for generating all other BiFC constructs, Birgit Kemmerling for BIR2-RFP and BIR3-CFP constructs, Katja Fröhlich for help with statistical analysis, Sarina Schulze for helpful discussions, Detlef Weigel for critical comments on the manuscript and Thorsten Nürnberger for critical comments and helpful discussions on the project.

## Notes

1 This work was supported by the University of Tübingen Graduate College “Of Plants and Men” and the Deutsche Forschungsgemeinschaft (SFB 766). E.S. was funded by the Max Planck Society. T.M. was funded by the Netherlands Organization for Scientific Research (867.15.020).

