## supplemental figures and tables for "The BIR2/BIR3-interacting Phospholipase D gamma 1 negatively regulates immunity in Arabidopsis"

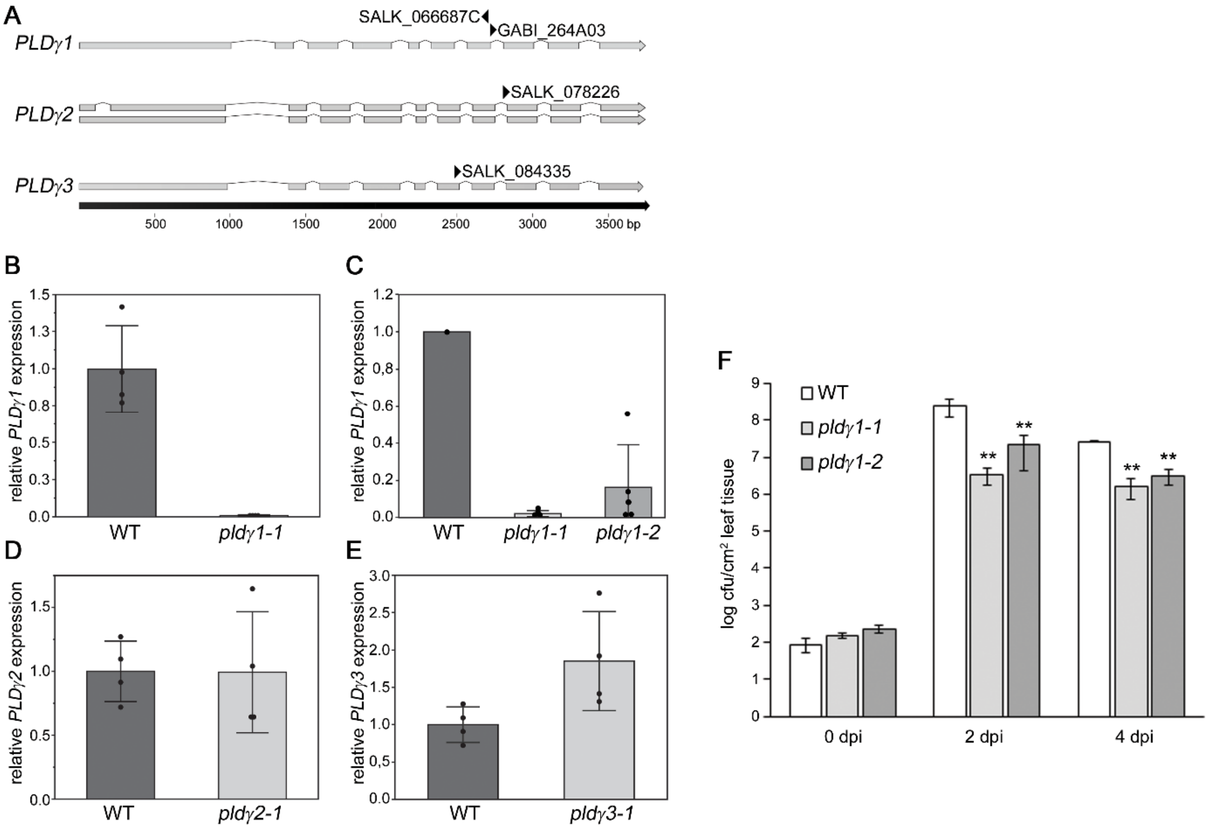


**Supplemental Figure S1.** Characterization of T-DNA insertion lines of *PLDγ* family members. A, Schematic display of the genes *PLDγ1*, *PLDγ2* and *PLDγ3* with exons indicated as grey boxes and introns as thin black lines. The black bold arrow at the bottom indicates nucleotide numbers in base pairs (bp). Position and names of T-DNA insertions are indicated with arrow heads above the respective gene. B-E, Transcriptional profiling of *PLDγ1* expression in the *pldγ1-1* mutant (B), of *PLDγ1* expression in the *pldγ1-1* line compared to *pldγ1-2* mutants (C), of *PLDγ2* expression in the *pldγ2-1* mutant (D) and of *PLDγ3* expression in the *pldγ3-1* mutant (E) as determined by RT-qPCR using total RNA prepared from leaf tissue of 6-week old Arabidopsis plants. Relative expression of the indicated gene was normalized to the level of the *EF-1α* transcript and calibrated to the level of the wild-type control, which was set to 1. Results are presented as the mean of 4 individual plants, error bars indicate SD (*n* = 4) and the whole experiment was repeated twice with similar results. F, Growth of *Pto* DC3000 in *pldγ1-1* and *pldγ1-2* mutants after infiltration of $10^4$ colony-forming units/ml. Data represent means ± SD of six replicate measurements per genotype per data point. Statistical significance compared with the wild type is indicated by asterisks (**p ≤ 0.05, Student's *t* test).

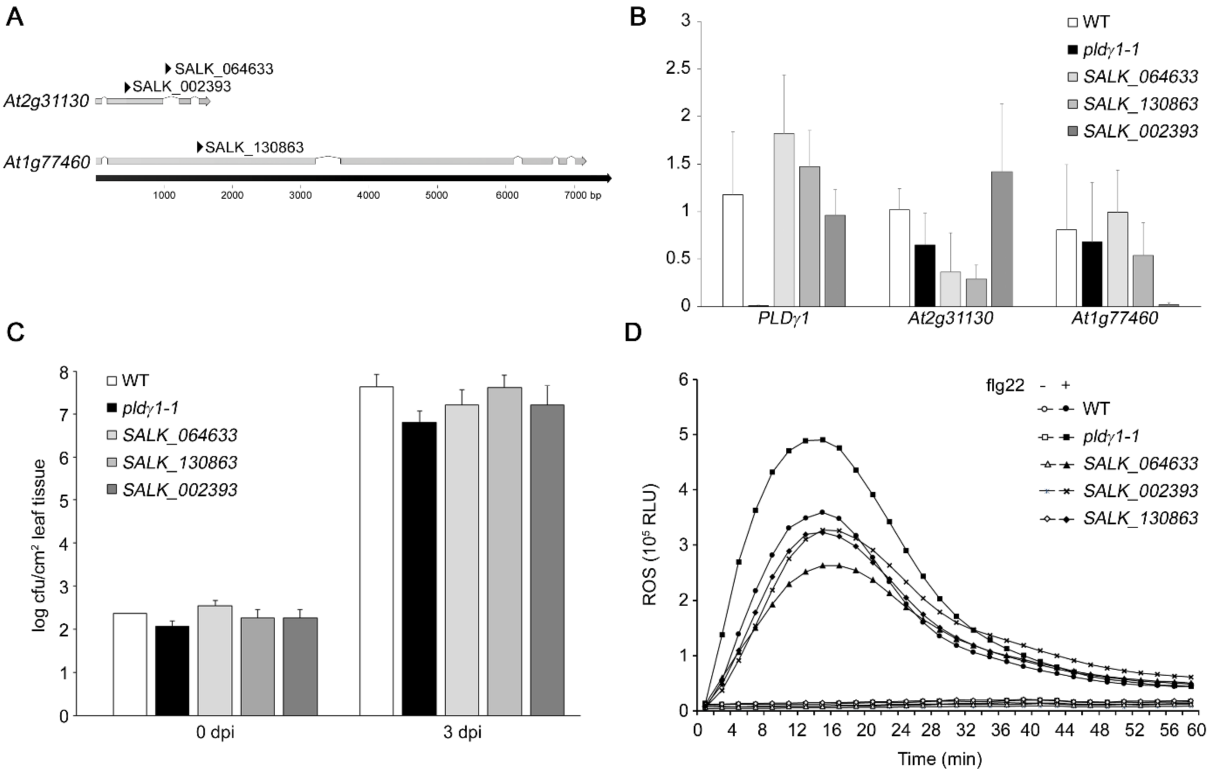


**Supplemental Figure S2.** Characterization of secondary mutations in the *pldγ1-1* mutant. A, Schematic display of the genes *At2g31130* and *At1g77460* with exons indicated as grey boxes and introns as thin black lines. The black bold arrow at the bottom indicates nucleotide numbers in base pairs (bp). Positions and names of the T-DNA insertions are indicated with arrow heads. B, Transcriptional profiling of gene expression of *PLDγ1, At2g31130* and *At1g77460* in the indicated mutant lines determined by RT-qPCR using total RNA prepared from leaf tissue of 6-week old Arabidopsis plants. Relative expression of the indicated gene was normalized to the level of the *EF-1α* transcript and calibrated to the level of the wild-type control, which was set to 1. Results are presented as the mean of 8 individual plants, error bars indicate SD ($n$ = 8) and the whole experiment was repeated twice with similar results. C, For bacterial infection, indicated mutant plants or the wild type as control were infiltrated with $10^4$ cfu/ml *Pto* DC3000. At day 0 and 3 post inoculation (dpi) bacterial growth was quantified by counting colony-forming units (cfu). The entire experiment was repeated three times with similar results. D, ROS accumulation after treatment of leaf pieces with or without 1 μM flg22 was measured over time in wild-type plants or indicated mutant lines and shown as relative light units (RLU). One out of four experiments is shown.

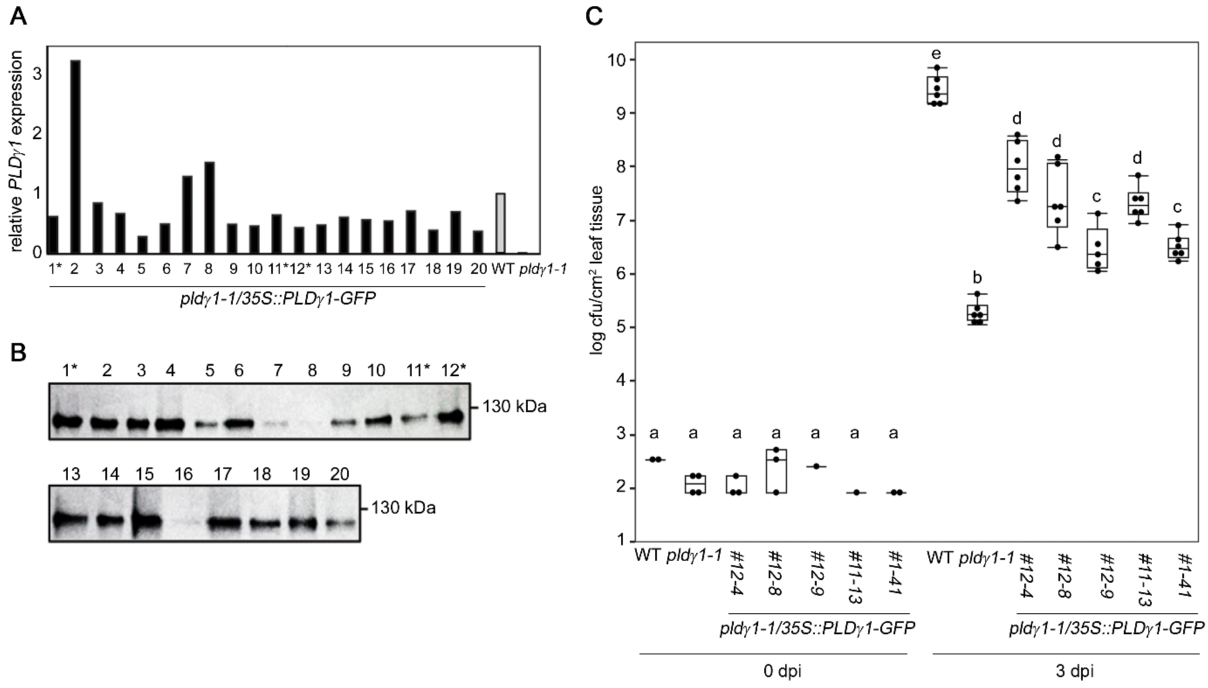


**Supplemental Figure S3.** Characterization of *pldγ1-1* complementation lines. Complementation lines were generated by stable transformation of the *pldγ1-1* mutant with a *35S::PLDγ1-GFP* construct. A, Restoration of *PLDγ1* transcript accumulation in the T1 generation. *PLDγ1* transcript levels were determined via RT-qPCR, normalized to *EF-1α* transcript levels and plotted as fold induction compared to wild-type plants as control, which were set to 1. Asterisks indicate mutant lines selected for further studies. B, Western Blot analysis of *PLDγ1-GFP* expression in the T1 generation. PLDγ1-GFP was immunoprecipitated from total protein extracts using Chromotek GFP-agarose beads and detected in Western Blot analysis using an anti-GFP antibody. Lines 1, 11 and 12 labelled with an asterisk were selected for further studies. C, Bacterial infection of *pldγ1-1* complementation lines. Wild type plants, *pldγ1-1* mutants or the T3 generation of complemented lines selected from (A) and (B) were infiltrated with $10^4$ cfu/ml of virulent *Pto* DC3000. At day 0 and 3 post inoculation (dpi) bacterial growth was quantified by counting colony-forming units (cfu). Box plots show minimum, first quartile, median, third quartile, and a maximum of log cfu/$cm^2$ leaf tissue (*n*=4 for 0 dpi; *n*=6 for 3 dpi). Labels a-d indicate homogenous groups according to post-hoc comparisons following one-way ANOVA (Tukey-Kramer multiple comparison analysis at a probability level of $p < 0.05$). The entire experiment was repeated with similar results.

A

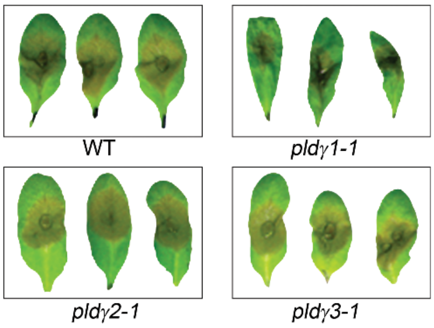


B

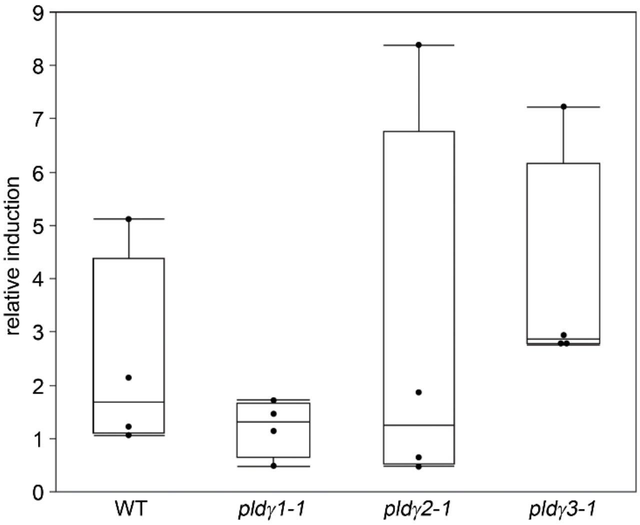


**Supplemental Figure S4.** *pldγ1* mutants are more resistant to *Botrytis cinerea* infection compared to *pldγ2* and *pldγ3* mutants*.* Leaves of 6-week old plants were inoculated with $5x10^6$/mL *Botrytis cinerea* spores. A, Three days after inoculation, symptom development was monitored and shown are three representative leaves per line as indicated. B, For the quantification of fungal biomass total DNA was extracted from infected leaf material and used for RT-qPCR. The relative amount of *B. cinerea* genomic *Actin*-DNA levels compared to Arabidopsis *Rubisco* (large subunit) levels was used to quantify fungal biomass. Box plots show minimum, first quartile, median, third quartile, and a maximum of fold induction of *Botrytis Actin* at day 3 compared to day 0 (*n*=4).

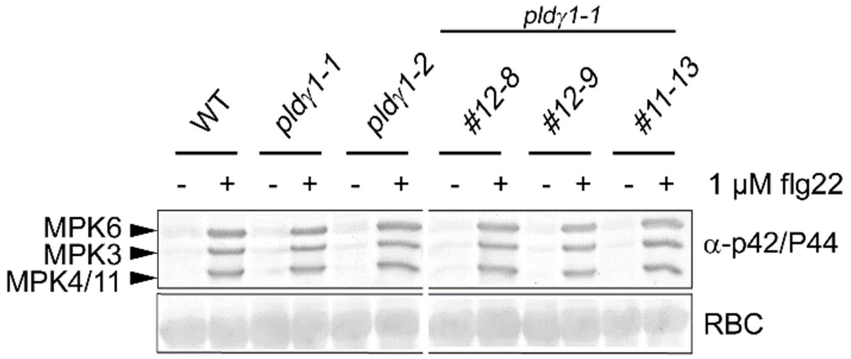


**Supplemental Figure S5.** Flg22-induced MAP kinase activation is not altered in *pldγ1* mutants or complementation lines. 10-day old seedlings of the wild type or the transgenic lines were treated for 15 min with 1 µM flg22 (+) or water as control (-). MAPK activation was visualized by Western Blot analysis using the phospho-p44/42 MAP kinase antibody. Ponceau S Red-staining of the membrane served as a loading control (RBC, Ribulose-bis-phosphate-carboxylase large subunit).

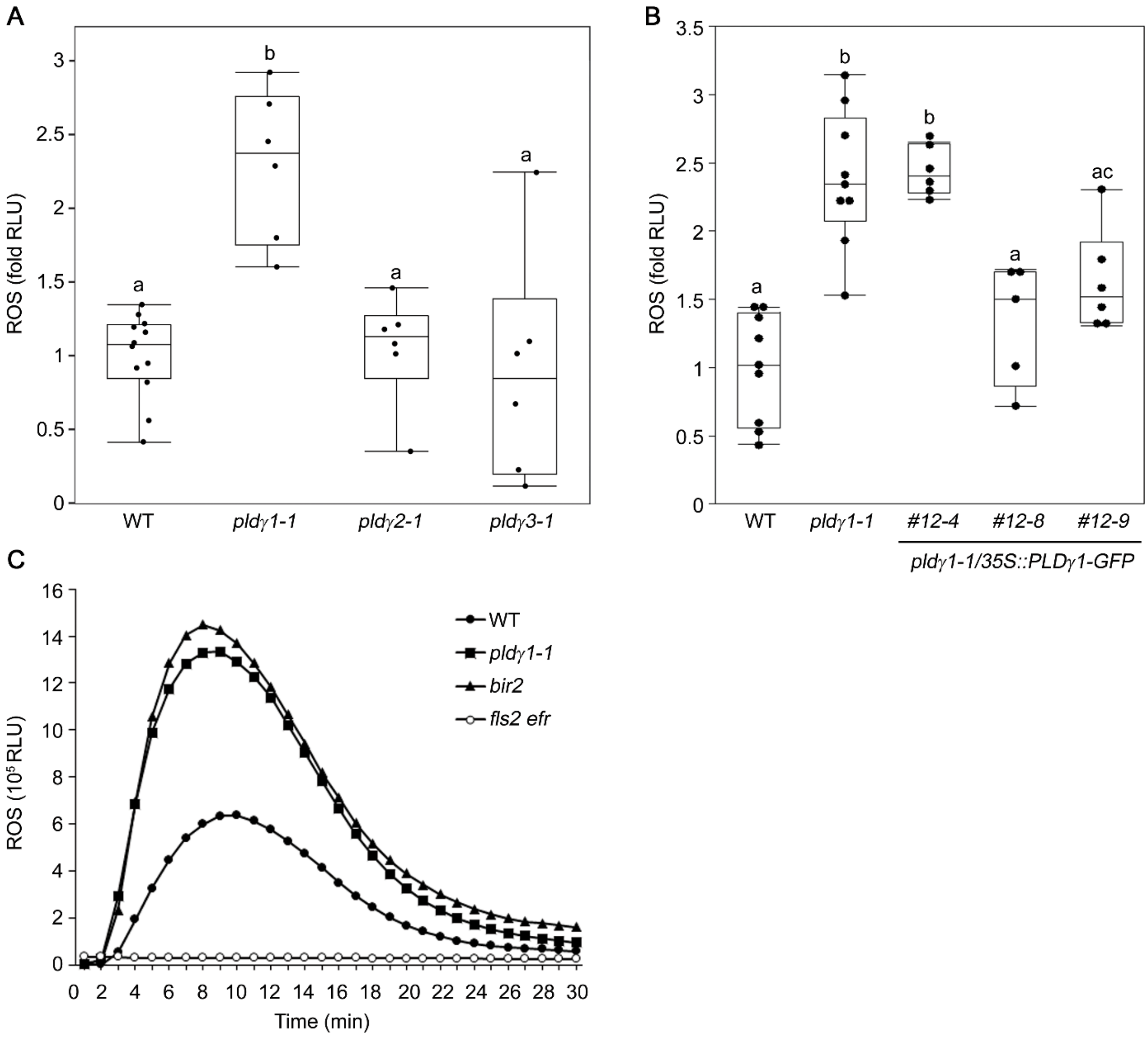


**Supplemental Figure S6**. ROS levels in *pldγ1-1* and *bir2* mutants are elevated compared to *pldγ2* and *pldγ3* mutant lines or *pldγ1-1* complementation lines. A and B, Arabidopsis leaf pieces of the indicated mutant or complementation line were treated with 1 µM flg22 or water as control, and ROS production was monitored over time. Shown are relative light units (RLU) expressed as fold induction to the mean of the wild-type, which was set to 1. Box plots show minimum, first quartile, median, third quartile, and a maximum of fold induction of peak value minus background value ($n \geq 6$). Water-treated samples had no detectable ROS production and are therefore not displayed in the figure. Labels a-c indicate homogenous groups according to post-hoc comparisons following one-way ANOVA (Tukey-Kramer multiple comparison analysis at a probability level of $p < 0.05$). C, ROS accumulation after treatment with 1 µM flg22 was measured over time in wild-type plants or *pldγ1-1*, *bir2* and *fls2 efr* mutant lines as described in (A, B) and shown as relative light units (RLU).

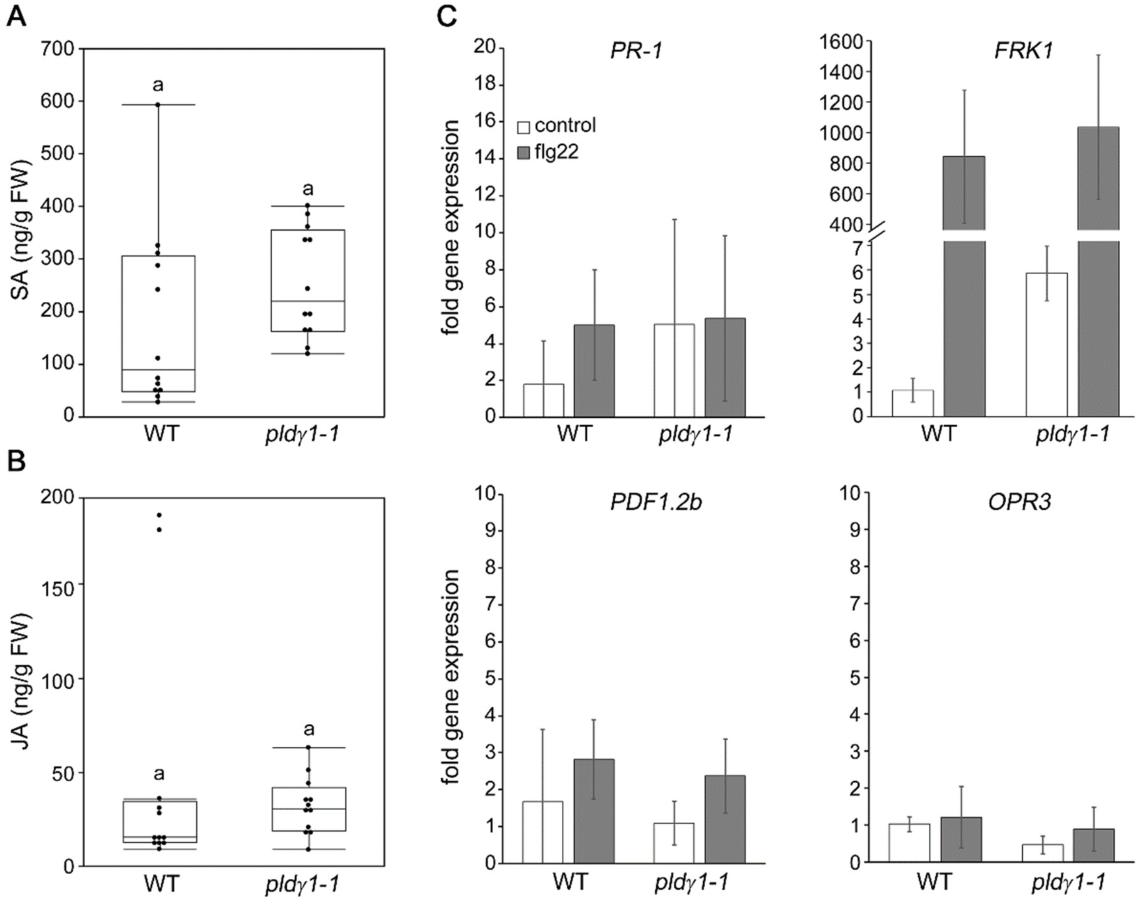


**Supplemental Figure S7.** PLDγ1 depletion does not affect SA and JA-signalling. A and B, Levels of salicylic acid (SA) (A) and jasmonic acid (JA) (B) were simultaneously determined from 200 mg leaf tissue of 8-week old wild-type (WT) plants or the *pldγ1-1* mutant. Box plots show minimum, first quartile, median, third quartile, and a maximum of the values represented in ng/g fresh weight (FW) ($n$=12). Label a indicates a homogenous group according to post-hoc comparisons following one-way ANOVA (Tukey-Kramer multiple comparison analysis at a probability level of $p < 0.05$). C, Transcriptional profiling of SA-and JA-marker genes by RT-qPCR. 9-day old seedlings of Arabidopsis wild type (WT) or the *pldγ1* mutant line were transferred from half-strength MS/1 % (w/v) sucrose plates into 500 µL liquid half-strength MS-medium (supplied with 1 % (w/v) sucrose) and treated with water (control) or 1 µM flg22 for 6 h. Relative expression of the indicated gene was normalized to the levels of the *EF-1α* transcript and calibrated to the levels of the control treatment in the wild-type seedlings, which was set to 1. Data present means of three technical replicates ± SD. All experiments shown were repeated at least three times with similar results.

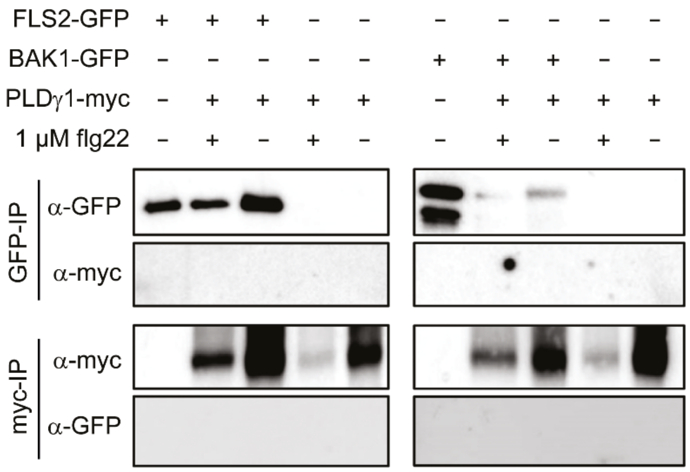


**Supplemental Figure S8.** PLDγ1 does not interact with FLS2 and BAK1. Western Blot analysis of transiently expressed proteins in *N. benthamiana* three days after infiltration. Leaf material was harvested 5 minutes after treatment with 1 µM flg22 (+) or water (-). After protein extraction each sample was split in two and was subjected to immunoprecipitation with either GFP- or myc-affinity beads as indicated. Immunoprecipitated and co-purified proteins were detected with anti-GFP or anti-myc antibodies as indicated. The experiment was repeated with similar results.

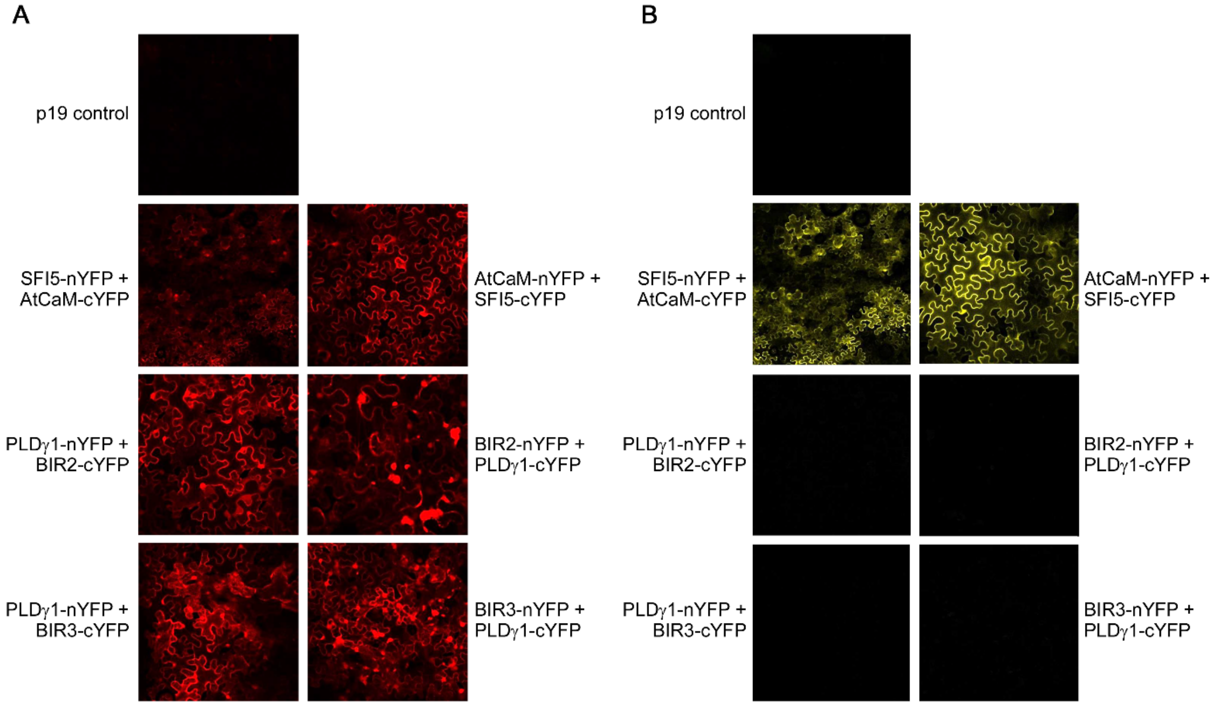


**Supplemental Figure S9.** PLDγ1 does not directly interact with BIR2 and BIR3 in BiFC. PLDγ1 C-terminally fused to either N- or C-terminal part of YFP was transiently expressed in *N. benthamiana* together with BIR2 or BIR3, fused to the respective other half of YFP using the pBIFCt2in1-CC vector system. A, Red fluorescence originating from the RFP-cassette present on the vector indicates successful transformation of *N. benthamiana* cells. B, Yellow fluorescence indicates reconstitution of functional YFP if fusion proteins interact. Interaction of Arabidopsis Calmodulin AtCaM1 and the *Phytophthora infestans* RXLR effector SFI5 was used as positive control. Fluorescence in epidermal cells was monitored 3 days after *Agrobacterium*-infiltration using confocal laser-scanning microscopy.

**Supplemental Table S1.** Primers used for genotyping T-DNA insertion lines.

| Gene name | Mutant name/ID | Primer name | Primer sequence (5‘ - 3‘) |
|---|---|---|---|
| *PLDγ1* (*At4g11850*) | *pldγ1-1* Salk_066687 | GT-PLDg1-LP | GGTGGGTTGCTAGTTTTTCG |
| | | GT-PLDg1-RP | CATCATGTTGCTATTCTCTGCTG |
| | *pldγ1-2* GK- 264A03 | GT-PLDg1-LP | GGTGGGTTGCTAGTTTTTCG |
| | | GT-PLDg1-RP | CATCATGTTGCTATTCTCTGCTG |
| *PLDγ2* (*At4g11830*) | *pldγ2* Salk_078226 | GT-PLDg2-LP | TGGAACTGGATGCCACTATTC |
| | | GT-PLDg2-RP | GGTTCCAACCTCTCTGTTTCC |
| *PLDγ3* (*At4g11840*) | *pldγ3* Salk_084335 | GT-PLDg3-LP | GGTTGTTTCAGTTGCATTTCA |
| | | GT-PLDg3-RP | GAACCCATTAAGGCAAAATCG |
| *At2g31130* | Salk_002393 Salk_064633 Salk_082136 | 31130-flCDSrv | TTACAGAAGCTCCCATTCAGATTC |
| | | 31130-flCDSfw | ATGGATTTTAAAGGTATAAAATGGGTTG |
| *At1g77460* | Salk_130863 | N630863-RP | CCTTTGACTTAGCATCAACGG |
| | | N630863-LP2 | ATCCAAGAACGGATCTTAGAG |
| SALK T-DNA | | LBb1.3 | ATTTTGCCGATTTCGGAAC |
| GABI-Kat T-DNA | | GABI-Kat LB | CCCATTTGGACGTGAATGTAGACAC |

**Supplemental Table S2.** Primers used in qPCR analyses.

| Gene name | Primer name | Primer sequence (5' – 3') |
|---|---|---|
| *At4g11850 (PLDγ1)* | qRT-PLDg1-fw1 | ACTTTTTCTGTCTTGGAACCAGAG |
| | qRT-PLDg1-rv1 | GCATTTGCATTTGCGTTTGGCTGA |
| *At4g11830 (PLDγ2)* | qRT-PLDg2-fw1 | TGCTCCCTTTGCGTCTAGGTTTCT |
| | qRT-PLDg2-rv1 | TCTATTCCAGCAGCAACACCACGA |
| *At4g11840 (PLDγ3)* | qRT-PLDg3-fw1 | GGTTTCCATAACACCCTTGTTGGT |
| | qRT-PLDg3-rv1 | TTCTCACCATCCACTTTATGATTCCT |
| *At2g31130* | qRT-AT2G31130-f | ATTGCAGAAAGGCAGAATCTGATG |
| | qRT-AT2G31130-r | GTCATTCTCCATCTTGTCAGGAAACAC |
| *At1g77460* | qRT2-77460-F | GTCGTGAGCAATAGCACTACTCC |
| | qRT2-77460-R | GCTATCTCAATGTCGAGTGATCTGG |
| *At2g19190 (FRK1)* | FRK1-100-F | AGCGGTCAGATTTCAACAGT |
| | FRK1-100-R | AAGACTATAAACATCACTCT |
| *At2g06050 (OPR3)* | At_OPR3_qF | GGCACCATGGTCTCTCCCGGAT |
| | At_OPR3_qR | ACCTCCCTTAGCGTGAACTGCTTC |
| *At2g14610 (PR-1)* | At_PR1_qF | GTGGGTTAGCGAGAAGGCTA |
| | At_PR1_qR | ACTTTGGCACATCCGAGTCT |
| *At2g26020 (PDF1.2b)* | PDF1.2b_qF | GGTACTTGGTCAGGAGTTTGC |
| | PDF1.2b_qF | ACTTGTGAGCTGGGAAGACA |
| *At1g07920/30/40 (EF-1α)* | EF1a-100-F | GAGGCAGACTGTTGCAGTCG |
| | EF1a-100-R | CACTTCGCACCCTTCTTGA |
| *Atcg00490 (Rubisco)* | AtRubisco-QF | GCAAGTGTTGGGTTCAAAGCTGGTG |
| | AtRubisco-QR | CCAGGTTGAGGAGTTACTCGGAATGCTG |
| *BC1G_08198 (Botrytis Actin)* | Bc_actin_qF | CCTCACGCCATTGCTCGTGT |
| | Bc_actin_qR | TTTCACGCTCGGCAGTGGTGG |
